# Genomic variation in educational attainment modifies Alzheimer’s disease risk

**DOI:** 10.1101/478776

**Authors:** Neha S. Raghavan, Badri Vardarajan, Richard Mayeux

## Abstract

**Objective:** To determine the putative protective relationship of educational attainment on Alzheimer’s disease (AD) risk using Mendelian randomization, and to test the hypothesis that by using genetic regions surrounding individually associated SNPs as the instrumental variable we can identify genes that contribute to the relationship.

**Methods:** We performed Mendelian randomization using genome-wide association study summary statistics from studies of educational attainment and AD in two stages. Our instrumental variable comprised of i) 1,271 SNPs significantly associated with educational attainment and ii) individual 2Mb regions surrounding the genome-wide significant SNPs.

**Results:** A causal inverse relationship between educational attainment and AD was identified by the 1,271 SNPs (odds ratio = 0.63; 95% CI, 0.54-0.74; p =4.08×10^−8^). Analysis of individual loci identified six regions that significantly replicated the causal relationship. Genes within these regions included *LRRC2*, *SSBP2*, and *NEGR1*; the latter a regulator of neuronal growth.

**Conclusions:** Educational attainment is an important protective factor for AD. Genomic regions that significantly paralleled the overall causal relationship contain genes expressed in neurons or involved in the regulation of neuronal development.

## Introduction

Education has consistently been identified as an important antecedent factor in Alzheimer’s disease (AD), whereby advanced educational attainment is thought to reduce AD risk^1^,^2^. In addition, educational attainment and AD may be genetically related based on genome-wide data showing a correlation of −0.31 (p=4×10^−4^)^3^. However, confounding factors that affect educational attainment such as socioeconomic status, nutrition and ethnicity blur the relationship with AD.

Mendelian randomization limits confounding by using instrumental variables associated with a risk factor to establish its effects on the outcome. The largest (n = 1,131,881) genome wide association study (GWAS) of educational attainment^4^ identified 1,271 significantly associated SNPs, and a separate 2013 GWAS from the International Genomics Alzheimer’s Project’s (IGAP) provide SNP association data to AD. Together, data from these published studies provided an opportunity to examine the role of educational attainment in AD through Mendelian randomization.

In GWAS analyses, the tag SNPs significantly associated with the phenotype may not be the causal SNP^5^. When pleiotropic effects and heterogeneity remain minimal, the instrumental variable composed of the index and all the associated SNPs in a region improves the validity of the relationship^6^, explains a greater percentage of variation of the phenotype and reduces heterogeneous effects of using fewer SNPs^7^. In addition to testing the overall hypothesis that educational attainment has a protective effect on AD risk, we tested the hypothesis that a subset of genetic loci related to educational attainment might individually significantly reflect the overall protective relationship with AD. In order to maximize detection, we included independent SNPs in a 2MB region surrounding each of the 1,271 SNPs significantly associated with educational attainment^3^.

## Methods

We used available summary statistics from the analysis of all discovery data excluding the 23andMe cohort in the largest GWAS meta-analysis of educational attainment^3^ (n = 766,345). Summary statistics from the IGAP GWAS of AD^8^, consisting of 54,162 individuals and 7,055,881 analyzed SNPs were used for the outcome. There is a small overlap of samples used from the educational attainment and AD GWAS. Sample overlap in a two-sample Mendelian randomization can cause bias in results; however, due to the large sample size of the educational attainment GWAS, and minimal overlap of sample, no appreciable bias was expected. The study population in both GWASs were of European descent.

### SNP selection

TwoSampleMR, an R package that performs Mendelian randomization using data from MR-Base^9^, was used in R (R v.3.3.1) to perform all analyses. Mendelian randomization was performed using two different schemes for selecting SNPs for the instrumental variable:

#### 1) 1,271 SNPs Analysis

The instrumental variables consisted of the 1,271 approximately independent SNPs (all with p < 5×10^−8^) significantly associated with educational attainment and were used to test for causality to AD. Independence for SNPs in the analysis was established using linkage disequilibrium (LD) clumping (r^2^≥0.001 within a 10,000 kb window).

#### 2) Individual locus analyses

Mendelian randomization was performed independently on each of the 1,271 loci using SNPs within 1MB upstream and downstream of individual SNPs. We used a more liberal inclusion criteria including SNPs associated with educational attainment at p<0.01 and clumped at r^2^ ≥0.1 within a 250kb window. Of the 1,271 independent loci, there were 243 regions, which contained more than one SNP within 2MB of another significant SNP. These were grouped together, resulting in 411 total independent locus tests.

### Mendelian randomization analyses

The strictest LD threshold was applied for the first analysis, using default settings in MR-Base; for analysis two we allowed for more SNPs with still minimal LD to be included in the analysis, as we hypothesized this would strengthen the per-loci analysis. Those SNPs remaining after LD clumping were queried within the IGAP summary statistics, if they were not present in that dataset, genetic proxies were found, and finally SNPs for which no proxies could be found were excluded. Next, SNPs were harmonized for the effect allele between the two GWAS datasets, or removed if harmonization predictions were inconclusive. For the joint 1,271 SNP analysis, 387 remain after LD clumping, 13 SNPs were neither found in IGAP nor had appropriate proxies and thus removed. Finally, 57 SNPs were removed due to low confidence that the effect allele of the exposure corresponded to the same allele in the outcome. Therefore, a total of 317 SNPs were included in the initial joint SNP analysis. For the regional analyses, SNPs were similarly removed if not found in IGAP or if allele harmonization failed. The inverse variance weighted (IVW) method was used to calculate odds ratios (OR). Results are reported as the OR (± 95% confidence interval) of AD risk per standard deviation increase in educational attainment in each test.

### Sensitivity analyses

Mendelian randomization requires that that the instrumental variables meets three requirements: it must be associated with the risk factor, not associated with any confounder of the risk factor or outcome, and is only associated with the outcome through the risk factor. Sensitivity analyses following the initial analysis provide confidence that assumptions of the instrumental variables were not broken. In addition to the IVW, the weighted median method was used to measure causality and provided consistent results with IVW when at least 50% of the instrumental variables were valid. Next, the intercept of the Mendelian Randomization-Egger test was used to determine potential horizontal pleiotropy, or an effect of the instrumental variables on a phenotype other than the outcome. As the intercept nears zero in the Mendelian randomization-Egger test, horizontal pleiotropy is reduced. The Steiger test of directionality was used to confirm directionality of the effect i.e. that the SNPs first affected educational attainment and subsequently that AD risk was affected through educational attainment. Finally, the radial regression analyses were performed for the inverse variance methods, which identifies any significant SNP outliers using the Cochran’s Q-statistic^10^.

## Results

Using the 1,271 SNPs associated with educational attainment, we detected statistically significant evidence for causation between educational attainment and AD, such that an increase of 4.2 years of educational attainment was associated with a 37% reduction in AD risk (OR - *scaled per SD*, *4.2 years* = 0.63; 95% CI, 0.54-0.74; p =4.08×10^−8^). In the per-loci analyses of the 1271 regions, two independent SNP regions demonstrated a statistically significant inverse relationship between educational attainment and AD risk (Bonferroni corrected threshold p<1.2 ×10^−4^). These regions include the neuronal growth regulator precursor (*NEGR1*) gene, leucine rich repeat containing 7 (*LRCC7*) gene, and prostaglandin e receptor 3 (*PTGER3*) gene (**Table 1**).

**Table 1.** Results from inverse variance weighted method and sensitivity analyses for Mendelian randomization analyses.

|  |  |  |  |  |  |  |  |
| --- | --- | --- | --- | --- | --- | --- | --- |
|  |  |  |  |  | Inverse variance weighted method | Sensitivity analysis |  |
| Top educational attainment SNPs (317 SNPs included) | | | | | Odds Ratio = 0.63; 95% CI, 0.54-0.74; $p = 4.08 \times 10^{-8}$ | Egger Regression: Intercept = 0.003; $p\text{-val} = 0.43$ | |
|  |  |  |  |  | Inverse variance weighted method |  | Sensitivity analysis |
| Locus | Chr | Gene(s) | Proposed role of gene(s) | # of SNPs | Odds Ratio (95% CI) | P-value | Egger Regression (intercept ± SE, $p\text{-val}$) |
| rs7552964<br>rs10789285<br>rs1024268<br>rs663251<br>rs481940<br>rs72677177<br>rs34122915<br>rs34305371<br>rs2568955<br>rs1445591<br>rs12028229<br>rs11210228<br>rs74091672<br>rs11210400<br>rs1569092<br>rs28482086 | 1 | *LRRC7/PTGER3/NEGR1/FPGT-TNNI3K/TNNI3K/TYW3/* | Necessary for synaptic spine architecture and function/ one receptor for prostaglandin E2/ Cell adhesion and, or regenerative axon sprouting/ read- through transcription/ MAPKKK family involved in cardiac physiology/ stabilizes codon-anticodon interactions | 89 | 0.31(0.19-0.50) | $1.92 \times 10^{-6}$ | .013 ± .006, $p\text{-val} = 0.05$ |
| rs2992632 | 5 | *SSBP2* | DNA damage response; maintenance of genome stability; telomere repair | 17 | 0.07(.02-.25) | $5.8 \times 10^{-5}$ | -.008 ± 0.017, $p\text{-val} = 0.66$ |

Sensitivity analyses were performed for all Mendelian randomization analyses. There was no indication of pleiotropy, reverse causality, or heterogeneity and weighted median method showed consistent results with IVW overall in the significant analyses.

## Discussion/Conclusions

Consistent with earlier reports, we found an inverse relationship of educational attainment with AD ^6^, ^11^, ^12^. In addition, we also identified regions that individually significantly replicated the causal relationship of education on AD^11^, ^12^, several of which were found to contain genes expressed in neurons or involved in regulation of neuronal development. These included regions surrounding a number of neuronally expressed genes. For example, one of the SNPs is within the leucine rich repeat containing 7 (LRCC7) gene which is crucial to dendritic spine architecture and function and may be involved in bipolar disorder^13^. Another SNP within the intronic region of the neuronal growth regulator precursor (*NEGR1*) gene is also highly expressed in neurons, and has been found to be associated with major depressive disorder^14^. Another SNP was found in the prostaglandin e receptor 3 (*PTGER3*) gene which is thought to be involved in the modulation of neurotransmitter release in neurons.

One limitation of using the Mendelian randomization method for outcomes with other strong non-genetic factors risk factors is that the confidence intervals tend to be large. However, this outcome is preferable to the bias that is inherent in epidemiological studies which unavoidably include confounders^15^. Taken together the results presented here, along with earlier reports, establish a putative causal relationship between educational attainment and AD.

The key next steps are to replicate these findings in diverse ethnic groups with more variable educational experiences, and to identify and validate specific variants within these loci that account for the association.

**Contributions**
All authors of the study: Neha Raghavan, Badri Vardarajan, Richard Mayeux, contributed to the study design, study analysis and writing and editing.

## Acknowledgements

Support for this work was provided by grants from the National Institute on Aging from the National Institutes of Health (U01AG032984 and RO1AG041797) and the National Center for Advancing Translational Sciences (TL1TR001875). We thank the International Genomics of Alzheimer’s Project and Lee et al. for providing summary statistics for this project.

## References

1. Caamano-Isorna F, Corral M, Montes-Martinez A, Takkouche B. Education and dementia: a meta-analytic study. Neuroepidemiology 2006;26:226–232.

2. Barnes DE, Yaffe K. The projected effect of risk factor reduction on Alzheimer’s disease prevalence. Lancet Neurol 2011;10:819–828.

3. Okbay A, Beauchamp JP, Fontana MA, et al. Genome-wide association study identifies 74 loci associated with educational attainment. Nature 2016;533:539–542.

4. Lee JJ, Wedow R, Okbay A, et al. Gene discovery and polygenic prediction from a genome-wide association study of educational attainment in 1.1 million individuals. Nat Genet 2018;50:1112–1121.

5. Schaid DJ, Chen W, Larson NB. From genome-wide associations to candidate causal variants by statistical fine-mapping. Nat Rev Genet 2018.

6. Burgess S, Bowden J, Fall T, Ingelsson E, Thompson SG. Sensitivity Analyses for Robust Causal Inference from Mendelian Randomization Analyses with Multiple Genetic Variants. Epidemiology 2017;28:30–42.

7. Pierce BL, Ahsan H, Vanderweele TJ. Power and instrument strength requirements for Mendelian randomization studies using multiple genetic variants. Int J Epidemiol 2011;40:740–752.

8. Lambert JC, Ibrahim-Verbaas CA, Harold D, et al. Meta-analysis of 74,046 individuals identifies 11 new susceptibility loci for Alzheimer’s disease. Nat Genet 2013;45:1452–1458.

9. Hemani G, Zheng J, Wade KH, et al. MR-Base: a platform for systematic causal inference across the phenome using billions of genetic associations. bioRxiv 2016. bioRxiv 078972; doi: https://doi.org/10.1101/078972

10. Bowden J, Spiller W, Del Greco MF, et al. Improving the visualization, interpretation and analysis of two-sample summary data Mendelian randomization via the Radial plot and Radial regression. Int J Epidemiol 2018;47:1264–1278.

11. E Anderson KW, G Hemani, J Bowden, R Korologou-Linden, GD Smith, Y Ben-Shlomo, LD Howe, E Stergiakouli. The Causal Effect Of Educational Attainment On Alzheimer’s Disease: A Two-Sample Mendelian Randomization Study. 2017. bioRxiv 127993; doi: https://doi.org/10.1101/127993

12. Larsson SC, Traylor M, Malik R, et al. Modifiable pathways in Alzheimer’s disease: Mendelian randomisation analysis. BMJ 2017;359:j5375.

13. Fiorentino A, Sharp SI, Kandaswamy R, Gurling HM, Bass NJ, McQuillin A. Genetic variant analysis of the putative regulatory regions of the LRRC7 gene in bipolar disorder. Psychiatr Genet 2016;26:99–100.

14. Hyde CL, Nagle MW, Tian C, et al. Identification of 15 genetic loci associated with risk of major depression in individuals of European descent. Nat Genet 2016;48:1031–1036.

15. Bautista LE, Smeeth L, Hingorani AD, Casas JP. Estimation of bias in nongenetic observational studies using “mendelian triangulation". Ann Epidemiol 2006;16:675–680.

